# Taxonomic identification accuracy from BOLD and GenBank databases using over a thousand insect barcodes from Colombia

**DOI:** 10.1101/2022.10.26.513939

**Authors:** Nathalie Baena-Bejarano, Catalina Reina, Diego Esteban Martínez-Revelo, Claudia A. Medina, Eduardo Tovar, Sandra Uribe-Soto, Jhon Cesar Neita-Moreno, Mailyn A. Gonzalez

**Author notes:** Corresponding authors (NBB), (MAG).

## Abstract

Recent declines of insect populations at high rates have resulted in the need to develop a quick method to determine their diversity and to process massive data for the identification of species of highly diverse groups. A short sequence of DNA from COI is widely used for insect identification by comparing it against sequences of known species. Repositories of sequences are available online with tools that facilitate matching of the sequences of interest to a known individual. However, the performance of these tools can differ. Here we aim to assess the accuracy in identification of insect taxonomic categories from two repositories, BOLD Systems, and GenBank. This was done by comparing the sequence matches between taxonomist identification and the suggested identification from the platforms. We used 1160 sequences in eight orders of insects from Colombia. After the comparison, we reanalyzed the results from a representative subset of the data from the subfamily Scarabaeinae (Coleoptera). Overall, BOLD systems outperformed GenBank, and the performance of both engines differed by orders and other taxonomic categories (species, genus and family). Higher rates of accurate identification were obtained at family and genus level. The accuracy was higher in BOLD for the orders Coleoptera at family, for Coleoptera and Lepidoptera at genus, and species level. The other orders performed similarly in both repositories. Moreover, the Scarabaeinae subset showed that in this group species were correctly identified when BOLD match percentage was above 93.4% and a total of 85% of the samples were correctly assigned to a taxonomic category. These results accentuate the great potential of the identification engines to place insects accurately into their respective taxonomic categories based on DNA barcodes and highlight the reliable use of BOLD Systems for insect identification in the absence of a large reference database for a highly diverse country.

## Introduction

More than a million species of plants and animals are in danger of extinction according to the latest global assessment of the intergovernmental platform on biological diversity and ecosystem services [1]. Declines in biodiversity and projections of a sixth mass extinction [2] have placed the impact of such losses on the agendas of governments all over the world [3]. Although insects constitute one of the most diverse groups on the planet, many species remain undiscovered and recent evidence suggests the decline of their populations at high rates [4,5]. To date, the risk of extinction of 12,161 species of insects has been evaluated according to the criteria of the IUCN red list and 2,291 species are considered “threatened” around the world [6].

Colombia is known as a biodiversity hotspot in the world. Although insects are the most studied class of animals in the country [7], research involving their genetic diversity is still currently underrepresented. Countries around the world are implementing massive sampling of insects with barcode generation under the premise that barcodes will allow us to assess biodiversity at a higher speed [8]. Barcodes in taxonomic works are showing an incredibly hidden diversity of insects with descriptions of hundreds of new species facilitated by DNA, morphology, and natural history [9,10]. As a megadiverse country, Colombia has great challenges to complete the inventory of its biodiversity and then the sustainable use of its resources. In particular, accelerating species identification using an integrative taxonomy is a priority.

Assessing biodiversity via sampling and DNA barcoding has been successfully applied in areas where coordinated sampling and barcoding facilitated the creation of extensive libraries for a particular region. An example of such collaboration is described in Morinière et al [8] where the Bavarian State Collection of Zoology (ZSM), the Barcoding Fauna Bavarica (BFB), and the German Barcode of Life (GBOL) project generated a library of 120,000 reliably identified species in Coleoptera, Diptera, Hymenoptera, and Lepidoptera for Germany and Central Europe. A closer example in the Neotropics is Costa Rica which has been leading the Bioalfa project and within the last 15 years has generated 45,000 insect species barcodes recorded [11]. Added to this initiative are other countries across the globe such as Canada, Ecuador, Sweden, and Singapore [9,12–16]. Moreover, the development and implementation of effective methods for species monitoring are important for assessing biodiversity, and overall health of ecosystems [8]. The species richness of insects in terrestrial ecosystems facilitates their use as bioindicators for measuring biodiversity and tracking the effects of changes in environmental conditions. The challenges faced by biodiversity declines have resulted in a need to generate new strategies for monitoring that are capable of generating vast amounts of data in a rapid and cost-effective manner. One such strategy that has been gaining traction in recent years has been the implementation of DNA barcoding.

Barcode sequences are generated from a standardized region that consists of short DNA sequences (generally between ∼300 to ∼700 bp). The mitochondrial Cytochrome c oxidase subunit I (COI) is most often used for insect barcoding. A DNA barcoding repository is built from sampling the DNA of individuals that are retained as vouchers, from which a photograph is taken. In the case of insects, most often a leg is removed from the specimen and used to obtain the DNA. In some cases, whole bodies of the samples are used when the genetic material available is limited by the size of the specimen. Although non-destructive DNA extraction methods are preferred to keep the morphology for future reference, photographs are suggested for facilitating identification before this step. This approach requires the creation of a large DNA reference database where the identification of unknown specimens is achieved through the information confirmed and curated in the database. The idea behind is that unknown samples could be identified by matching their sequences to the species that are curated from the database. The Barcode of Life Data Systems BOLD (http://www.boldsystems.org) [17] is a platform specifically developed for this type of information. The BOLD platform provides the identifications match based on BINs (Barcode Index Numbers) [18] that are automatically generated through algorithms for the specimens. Although unidentified species coded in BINs can help estimating diversity, the identification of species is also possible using the Basic Local Alignment Tool (BLAST) from GenBank [19,20]. BLAST allows matching sequences to a query sequence. The search consists of comparing nucleotide sequences hosted in the database and to provide statistical significance of these matches. The main goal from BLAST is not necessarily to provide species identification because the same principle can be used for the search of gene families; uses of identification databases are vast in biology. Scientists frequently use these two platforms to identify species from barcodes. An assessment of these two main public repositories from curated samples across taxa found a slightly better performance from GenBank vs BOLD for insects (n=17) at species and genus level identification without statistical significance, and more even results between repositories for plants (n=61) and macro-fungi (n=16) [21]. Later, a revision of this insect data found the results to be more similar than previously reported for this group [22]. Accurate identification of species and overall composition in ecosystems based on sampling is a key factor for conservation initiatives. Therefore, evaluating the performance of these platforms assigning specimens to a specific taxonomic category level has a significant impact on the advancement of other fields.

Here, we analyze a large inventory of over 1,000 insect barcodes for Colombia to the database which includes the first strepsipteran barcode from Colombia. We aim to compare the matches of the taxonomist identification to species, genus, and family level with the search engines of BOLD and BLAST service from GenBank in order to provide suggestions and next steps for the use of these tools. We assess the accuracy of the species identification engine from BOLD by using a subset of insects corresponding to Scarabaeinae subfamily (Coleoptera: Scarabaeidae).

## Materials and Methods

### Collection Locality/Geography

Insect samples were collected from ten departments of Colombia through different projects (Colombia BIO, Santander BIO). A majority of the samples (81.0%) came from three departments of Colombia (Antioquia, Vichada, and Santander), the spatial data of the localities were processed in ArcGIS 10.2 (License E300 04/26/2013) (Fig 1). The elevation of the samples covered a wide range with the lowest sampling occurring at 56 masl and the highest at 3563 masl. Although samples were obtained with seven different entomological collecting methods (malaise, pitfall, light trap, hand-picked, entomological net, Van Someren-Rydon trap and folding net), nine malaise traps accounted for 62.6% of the total collected material, followed by pitfall 17.2%, and light trap 8.1%.

**Fig 1.**
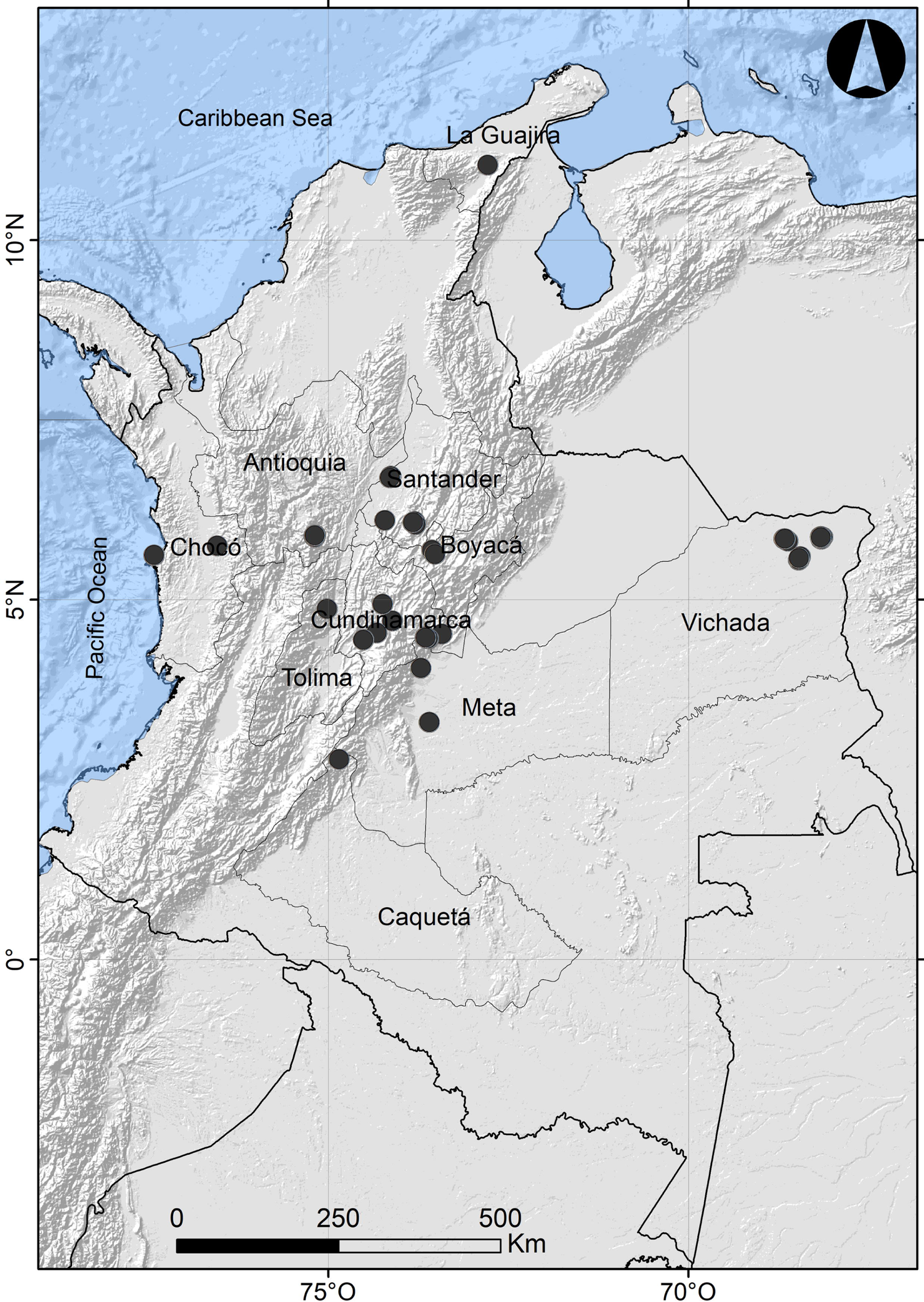
Map of Colombia showing collecting localities in black dots. Marine polygon, department boundaries, and physical labels were obtained from Natural Earth (http://www.naturalearthdata.com/). Hillshade map based on SRTM30_PLUS v8 [23].

### Taxa description

This dataset includes insects from the orders Coleoptera, Diptera, Hymenoptera, Lepidoptera, Hemiptera, Odonata, Psocodea and Strepsiptera, from which Coleoptera is the most represented order in the dataset (35%). The specimens were identified from morphology by students working on their thesis and by specialists. As a result of varying expertise, we provide an identification that goes from order to species level (S1 Table); additionally in Scarabaeinae, identification was done to morphospecies level following González-Alvarado et al [24]. Currently, few samples (7%) have been identified to lower levels from molecular data by BOLD Data Manager. The specimens were deposited at collections with 999 samples between the Entomological and Tissue collection of the Institute Alexander von Humboldt (IAvH-E and IAvH-CT), Museo Javeriano de Historia Natural (MPUJ) at Pontificia Universidad Javeriana with 63, Museo Francisco Luis Gallego (MEFLG) at Universidad Nacional de Colombia, Sede Medellin with 55, and Colección Entomológica Universidad de Antioquia (CEUA) with the remaining 43 samples. Records of some samples are also available for search in SIB Colombia.

### DNA sampling and extraction

One leg was removed from each insect, when possible, but some samples required the use of the full body to provide enough DNA. All samples were sent to the molecular laboratory of Instituto Humboldt at CIAT, Palmira. Extraction protocols were conducted following the high throughput DNA isolation from Ivanova et al [25]. The concentration of selected DNA samples was estimated by quantification using the NanoDrop 1000 (Thermo Fisher Scientific). Amplification of cytochrome c oxidase I (COI) fragments (658 bp) took place by polymerase chain reaction (PCR) with a laboratory prepared master mixture which contained 2 μL of template DNA (∼10-50 ng), 1X Taq buffer ((NH4)2SO4), 200 μM of each deoxynucleoside triphosphate, 2 mM MgCl2, 0.2 μM of each primer, 0.4 μg/μL bovine serum albumin, and 1 U of Taq DNA polymerase. A cocktail of four primers was used: LCO1490 5’-GGTCAACAAATCATAAAGATATTGG-3’, HCO2198 5’-TAAACTTCAGGGTGACCAAAAAATCA-3’ [26], Lep-F1 5’-ATTCAACCAATCATAAAGATATTGG-3’ and Lep-R1 5’-TAAACTTCTGGATGTCCAAAAAATCA-3’ [27]. Primers were selected following Canadian Centre for DNA Barcoding (CCDB) suggested primers for taxonomic groups. The PCR cycling conditions consisted of a first cycle of denaturation at 94 °C for 3 min; followed by 5 cycles of 94 °C for 30 s, 45 °C for 40 s, and 72 °C for 1 min; then 35 cycles of 94 °C for 30 s, 51 °C for 40 s, and 72 °C for 1 min; and a final extension cycle at 72 °C for 5 min. PCR products were visualized by electrophoresis on 1.5% agarose gels stained with SYBR Safe (Thermo Fisher Scientific) and using 1X TBE. The ExoSAP-IT protocol was used to clean PCR products, after which Sanger sequencing of amplicons were carried out at The Universidad Nacional de Colombia and the University of the Andes. Sequences were assembled and edited in Geneious 10.0.9 2017 (Auckland, New Zealand) [28]. All files were uploaded in BOLD systems and are found in the dataset “insects-all” (DS-CBIH2020; Dataset available at: DOI will be made available after acceptance).

### Data analysis

#### BOLD Batch ID engine

To identify samples to species level, we ran BOLD’s Identification Engine in datasets created for insect orders in July 2020. The identification engine allows the user to select the search within the COI Species Database or COI Full Database. We selected COI Species Database which will match the sequence against an identified species as opposed to individuals without identification to species level. We kept filters as default for a minimum overlap of 300 bp and sequence length ≥ 100 bp. We ran the analysis with 97% similarity, a percentage that accounts for within species variation and error of the sequence [8]. Under the criteria selected of 97% numerous species ID were not obtained. We decided to lower the criteria and include matches using the 80% similarity filter due to the lack of matches at 97%. The 80% similarity is the minimum filter allowed in BOLD. A match was accepted as correct when the identification by the taxonomist was in agreement by the suggested identification by BOLD. The ID search engine provides a list of maximum 100 closest specimens, and we selected the first suggestion from the list that has the highest percentage match and base pair overlap. We included an alternative match when the suggestion was done towards a sequence from our project and ran statistical tests separately to prevent circular analysis of results. Matches were checked at species, genus, and family level.

#### BLAST

We used BLAST from GenBank in Geneious 10.0.9 2017 by default to look for matches of the sequences. The parameters included were the database search from Nucleotide collection, program set as the MegaBlast, results displayed as a Hit table, and retrieve Matching region with Maximum hits of 100. The search was set to provide a list of 100 closest nucleotide sequences, from which we selected the first suggestion that obtained the highest Bit-score or Max score (name when the BLAST search is done from the GenBank website). Most of the time the E value was 0 (or closest to zero) and the Query cover 100%. Matches were checked at species, genus and family level.

The identification analyses in both approaches (BOLD and Blast) followed the assumption that species from the databases are correctly identified; however, this approach might not be accurate in cases where the most similar species were wrongly identified from the databases.

### Dung beetles (Scarabaeinae) case study

The Scarabaeinae subfamily was selected due to the high representation of this group in the samples of Coleoptera (42%) and from the total insect samples (15%). In order to evaluate the accuracy of species identification suggested by BOLD, we consulted taxonomists with expertise in the identification of the Scarabaeidae family. They were asked to reanalyzed the results and mention if they agree with the species and genus identification suggested by BOLD and to explain why, as follows: Correctly Identified Genus, Correctly Identified Species, Probable Correctly Identified Species, Incorrect Identification (Table 1). When available, specimens deposited at the IAVH-E collection and photographs uploaded to BOLD were checked for specimens to corroborate the taxonomy, as field notes from the collectors.

**Table 1.** List of categories used by the taxonomist to define the identification results suggested by BOLD with the Scarabaeinae subfamily.

| Category | Definition |
| --- | --- |
| Correctly identified genus | Agreement between the taxonomist and BOLD at the genus level |
| Correctly identified species | Agreement between the taxonomist and BOLD at the species level |
| Probable correctly identified species | Agreement between the taxonomist morphospecies and the species suggested by BOLD (the genus matched and probable species) |
| Incorrect identification | Disagreement between the taxonomist and BOLD (the genus or species did not match) |

### Statistical analysis

Chi-square analyses were performed between match results for BOLD and GenBank at the studied taxonomic levels (species, genera and family) to evaluate if there was a significant difference between the platforms performance. An additional series of Chi-square analyses were performed to identify differences in the platforms by each order of insect at the species, genus, and family level. We adjusted our significance threshold at each taxonomic level based on the number of orders compared using the Bonferroni Correction. Flagged records were removed from the analysis.

The non-parametric Kruskal-Wallis test in PAST, Version 4.05 [29] was used to determine any significant differences of medians for BOLD % match of the correct Scarabaeinae categories against incorrect identification. Assumptions for an ANOVA were not fulfilled. Dunn’s post hoc was carried out to identify differences between categories.

## Results

### Description of the data

A total of 1160 sequences of insects from Colombia were generated since 2019. From this set, 1088 sequences were barcode compliant and placed in 708 BINS (Table 2). Of these 708 BINS, 500 were identified as unique (Fig 2). The most frequent BINS were all in Coleoptera, BOLD:ADN3981 *Paulosawaya* sp. (11 specimens), BOLD:ADO6609 *Paulosawaya ursina* (Blanchard, 1850) (10 specimens), and BOLD:ADJ9394 Ptilodactylidae (9 specimens). From the specimens with barcode compliant 33 were flagged as problematic records in most instances due to suspected contamination, specimen mix-up or misidentification.

**Table 2.** Summary of specimens and barcodes data by orders after removal of flagged records.

| Order | Specimens<br># | Flagged<br>record | Family<br>count | Specimen<br>count<br>IDed<br>Family | Unknown | Genera<br>count | Specimen<br>count<br>IDed<br>Genus | Species<br>count | Specimen<br>count IDed<br>Species | Barcode<br>compliant | Species<br>count | BINS | Unique<br>Bins | Non-Unique<br>Bins |
| --- | --- | --- | --- | --- | --- | --- | --- | --- | --- | --- | --- | --- | --- | --- |
| Coleoptera | 411 | 14 | 25 | 397 | 0 | 57 | 271 | 63 | 117 | 369 | 57 | 218 | 184 | 34 |
| Diptera | 277 | 14 | 24 | 257 | 6 | 4 | 22 | 0 | N/A | 257 | 0 | 165 | 116 | 49 |
| Hemiptera | 82 | 4 | 11 | 78 | 0 | 0 | N/A | 0 | N/A | 68 | 0 | 58 | 56 | 2 |
| Hymenoptera | 201 | 2 | 10 | 199 | 0 | 16 | 47 | 2 | 3 | 176 | 2 | 142 | 112 | 30 |
| Lepidoptera | 166 | 0 | 8 | 161 | 5 | 57 | 154 | 79 | 128 | 163 | 73 | 104 | 18 | 86 |
| Odonata | 17 | 0 | 6 | 17 | 0 | 14 | 17 | 13 | 15 | 16 | 13 | 15 | 8 | 7 |
| Psocodea | 5 | 0 | 4 | 5 | 0 | 0 | N/A | N/A | N/A | 5 | N/A | 5 | 5 | 0 |
| Strepsiptera | 1 | 0 | 1 | 1 | 0 | 1 | 1 | 0 | N/A | 1 | 0 | 1 | 1 | 0 |
| **Total** | **1160** | **34** | **89** | **1115** | **11** | **149** | **512** | **157** | **263** | **1055** | **145** | **708** | **500** | **208** |

**Fig 2.**
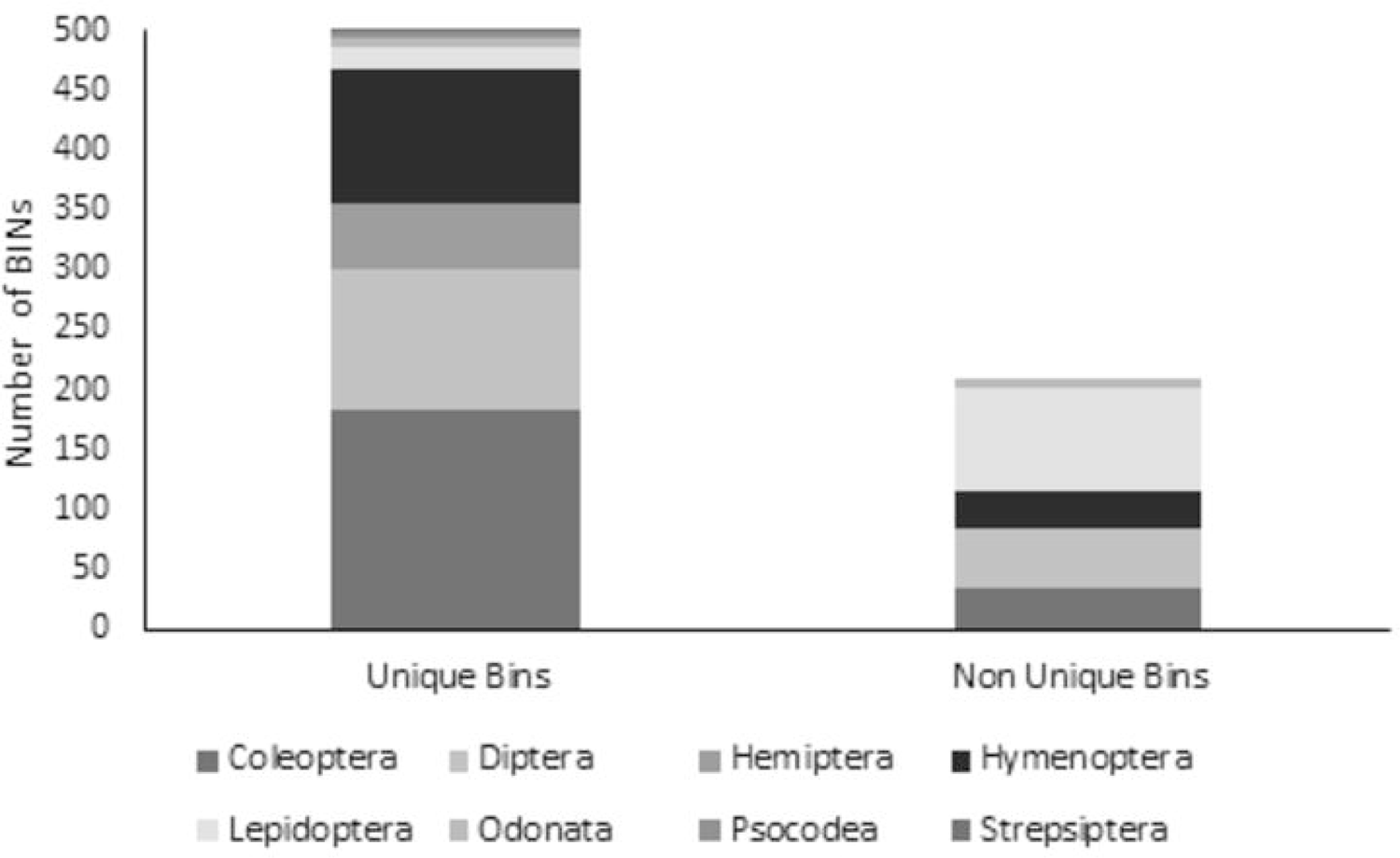
Summary of Barcode index numbers (BINS) obtained by orders of insects. A total of 500 BINs were identified as unique and 208 BINs as non-unique in the platform BOLD Systems.

### Coleoptera

S1 Table and S2 Table; Barcodes were obtained from 369 out of 411 coleopteran specimens and placed in 218 BINS, from which 184 (84.4%) were identified as unique in BOLD. The samples were assigned to 25 families, 57 genera and 63 species.

BOLD search engine: The search at 80% provided identifications for a total of 395 specimens. A match in agreement with the taxonomist was found for 337 (85%) specimens at the family level. For genus, the search resulted in 189 (70%) specimens of 271 and for species in 57 (49%) specimens of 117 total.

BLAST: The search provided identifications for a total of 379 specimens. A match in agreement with the taxonomist was found for 294 (78%) specimens at the family level. For genus, the search resulted in 83 (33%) specimens of 255 and for species in 20 (20%) specimens of 101 total.

### Diptera

S1 Table and S2 Table; Barcodes were obtained from 257 out of 277 dipteran specimens and placed in 165 BINS, from which 116 (70.3%) were identified as unique in BOLD. The samples were assigned to 24 families and 4 genera. There were 6 specimens not assigned to any category.

BOLD search engine: The search at 80% provided identifications for a total of 245 specimens. A match in agreement with the taxonomist was found for 202 (82%) specimens at the family level, and for genus, 16 of 18 specimens (89%) matched the identification of the taxonomist. No species were identified by the taxonomist.

BLAST: The search provided identifications for a total of 241 specimens. A match in agreement with the taxonomist was found for 184 (76%) specimens at the family level, and for genus 13 of 14 specimens (93%) matched the identification of the taxonomist. No species were identified by the taxonomist.

### Hemiptera

S1 Table and S2 Table; Barcodes were obtained from 68 out of 82 hemipteran specimens and placed in 58 BINS, from which 56 (97%) were identified as unique in BOLD. The samples were assigned to 11 families.

BOLD search engine: The search at 80% provided identifications for a total of 77 specimens. A match in agreement with the taxonomist was found for 66 (86%) specimens at the family level. No genera or species were identified by the taxonomist.

BLAST: The search provided identifications for a total of 67 specimens. A match in agreement with the taxonomist was found for 55 (82%) specimens at the family level. No genera or species were identified by the taxonomist.

### Hymenoptera

S1 Table and S2 Table; Barcodes were obtained from 176 out of 201 hymenopteran specimens and placed in 142 BINS, from which 112 (78.9%) were identified as unique in BOLD. The samples were assigned to 10 families, 16 genera and 2 species.

BOLD search engine: The search at 80% provided identifications for a total of 199 specimens. A match in agreement with the taxonomist was found for 193 (97%) specimens at the family level. At the genus level, the search resulted in 46 (98%) of 47 specimens and 3 of 3 (100%) specimens at the species level.

BLAST: The search provided identifications for a total of 182 specimens. A match in agreement with the taxonomist was found for 176 (97%) specimens at the family level. At the genus level, the search resulted in 38 (97%) of 39 specimens and 0 of 3 specimens at the species level.

### Lepidoptera

S1 Table and S2 Table; Barcodes were obtained from 163 out of 166 lepidopteran specimens and placed in 104 BINS, from which 18 (17%) were identified as unique in BOLD. The samples were assigned to 8 families, 57 genera and 79 species, 5 not assigned to any category.

BOLD search engine: The search at 80% provided identifications for a total of 161 specimens. A match in agreement with the taxonomist was found for 147 (91%) specimens at the family level. For genus, the search resulted in 140 (91%) specimens of 154 and for species in 86 (68%) specimens of 127 total.

BLAST: The search provided identifications for a total of 154 specimens. A match in agreement with the taxonomist was found for 143 (93%) specimens at the family level. For genus, the search resulted in 117 (79%) specimens of 148 and for species in 49 (49%) specimens of 99 total.

### Odonata

S1 Table and S2 Table; Barcodes were obtained from 16 out of 17 Odonata specimens and placed in 15 BINS, from which 8 (53.3%) were identified as unique in BOLD. The samples were assigned to 6 families, 14 genera and 13 species.

BOLD search engine: The search at 80% provided identifications for a total of 17 specimens. A match in agreement with the taxonomist was found for 17 (100%) specimens at the family level. For genus, the search resulted in 13 (76%) specimens of 17 and for species in 6 (46%) specimens of 13 total.

BLAST: The search provided identifications for a total of 17 specimens. A match in agreement with the taxonomist was found for 17 (100%) specimens at the family level. For genus, the search resulted in 9 (56%) specimens of 16 and for species in 3 (20%) specimens of 15 total.

### Psocodea

S1 Table and S2 Table; Barcodes were obtained from 5 out of 5 psocodean specimens and placed in 5 BINS, each identified as unique in BOLD (100%). The samples were assigned to 4 families.

BOLD search engine: The search at 80% provided identifications for a total of 5 specimens. A match in agreement with the taxonomist was found for 2 (40%) specimens at the family level. No genera or species were identified by the taxonomist.

BLAST: The search provided identifications for a total of 4 specimens. A match in agreement with the taxonomist was found for 1 (25%) specimen at the family level. No genera or species were identified by the taxonomist.

### Strepsiptera

S1 Table and S2 Table; A barcode was obtained from one male specimen and placed in 1 new BIN. The sample was identified to the genus *Caenocholax*.

BOLD search engine: The search at 80% provided identification for the 1 specimen. A match in agreement with the taxonomist was found for it (100%) at the family and genus level. No species were identified by the taxonomist.

BLAST: The search provided identification for the 1 specimen. A match in agreement with the taxonomist was found for it (100%) at the family and genus level. No species were identified by the taxonomist.

### Performance of BOLD and Blast (GenBank) identification based on taxonomist match

The performance of BOLD vs BLAST for identification of specimens was significantly greater with BOLD for the pooled comparison of all species (x2 = 29.5, df = 1, P < 0.001), genera (x2 = 67.7, df = 1, P < 0.001), and families (x2 = 8.3, df = 1, P = 0.004). Comparisons of the performance of BOLD and BLAST by Orders at the different taxonomic levels revealed significantly more matches with BOLD (Fig 3), at the family level (Fig 3A) in Coleoptera (x2 = 7.7, df = 1, P = 0.006), at genus level (Fig 3B) in Coleoptera (x2 = 72.8, df = 1, P < 0.001) and Lepidoptera (x2 = 8.4, df = 1, P = 0.004) and at the species level (Fig 3C) in Coleoptera (x2 = 19.8, df = 1, P < 0.001) and Lepidoptera (x2 = 7.7, df = 1, P = 0.006). No differences were found for the other orders tested at the different taxonomic categories. Same results were found when analyses were performed with matches to data from our project, the only comparison that changed was at the genus level where no significant differences were found for Lepidoptera (S3 Table).

**Fig 3.**
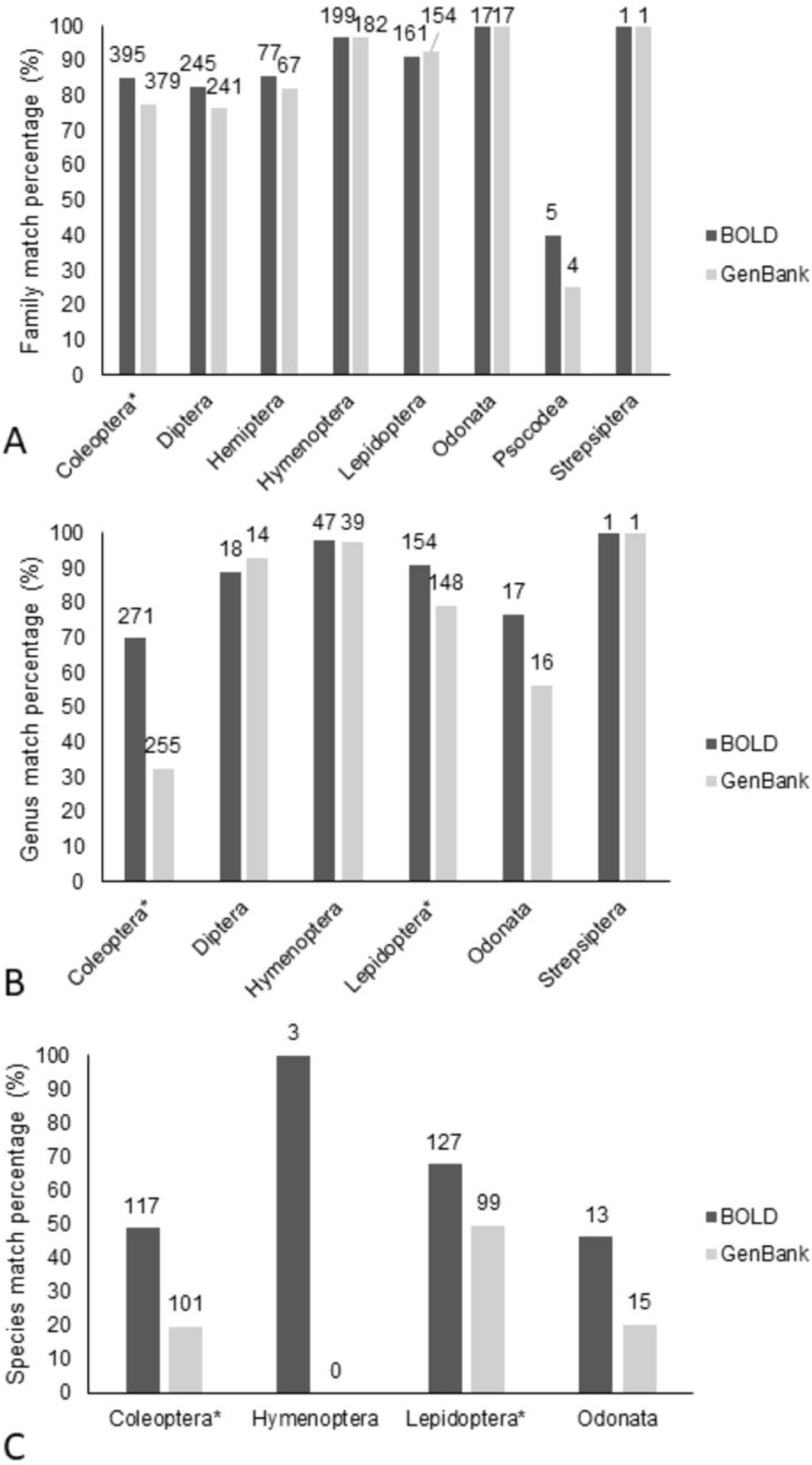
Comparison by order of performance of BOLD Systems and BLAST for identification of specimens in percentages. Correct matches for BOLD Batch ID engine at 80% are shown in dark gray. Correct matches from MegaBlast search by default (GenBank) are shown in light gray. *Is next to orders where Chi-square analyzed with Bonferroni correction was significant. The sample size is displayed on top of each column. A) by families. B) by genera C) by species.

### Case of study Scarabaeinae subfamily (Coleoptera: Scarabaeidae)

A total of 172 specimens were analyzed. The samples in the subfamily Scarabaeinae were assigned to 16 genera and 27 species (S1 Table).

BOLD search engine: The search at 80% provided identifications for the 172 specimens (Fig 4A, S2 Table). A match in agreement with the taxonomist was found for 82 (80%) specimens of 103 at the genus level. For species, the search resulted in 55 (78%) specimens of 69 total. Based on this information, these records were classified as 137 (80%) in agreement with the taxonomist and 35 (20%) in disagreement (Table 3, Fig 4A).

**Table 3.** Summary of matches between the taxonomist and BOLD identifications for the subfamily of dung beetles Scarabaeinae. The first section (I) corresponds to the initial matches performed in agreement with the taxonomist after the search engine in BOLD at 80%. The second section (II) in grey includes the categories used by the taxonomist to reevaluate the identifications provided by BOLD in (I) and the final matches in agreement.

|  |  |  |  | Matches<br>BOLD &<br>Taxonomist | Count | Total | % | Difference %<br>(disagreement<br>with<br>taxonomist) |
| --- | --- | --- | --- | --- | --- | --- | --- | --- |
| (I) BOLD search<br>engine<br>identifications |  |  |  | Genus | 82 | 103 | 80 |  |
|  |  |  |  | Species | 55 | 69 | 78 |  |
|  |  |  |  | **Sum**<br>**agreement** | **137** | 172 | **80** | 20 |
| Categories |  | Count | % |  |  |  |  |  |
| (II) Taxonomist<br>corroborated<br>identifications –<br>*de novo* | Correctly identified<br>genus | 52 | 30 | Genus | 52 | 73 | 71 |  |
|  | Correctly identified<br>species | 77 | 45 | Species | 95 | 99 | 96 |  |
|  | Probable correctly<br>identified species | 18 | 10 |  |  |  |  |  |
|  | Incorrect identification | 25 | 15 | **Sum**<br>**agreement** | **147** | 172 | **85** | 15 |

**Fig 4.**
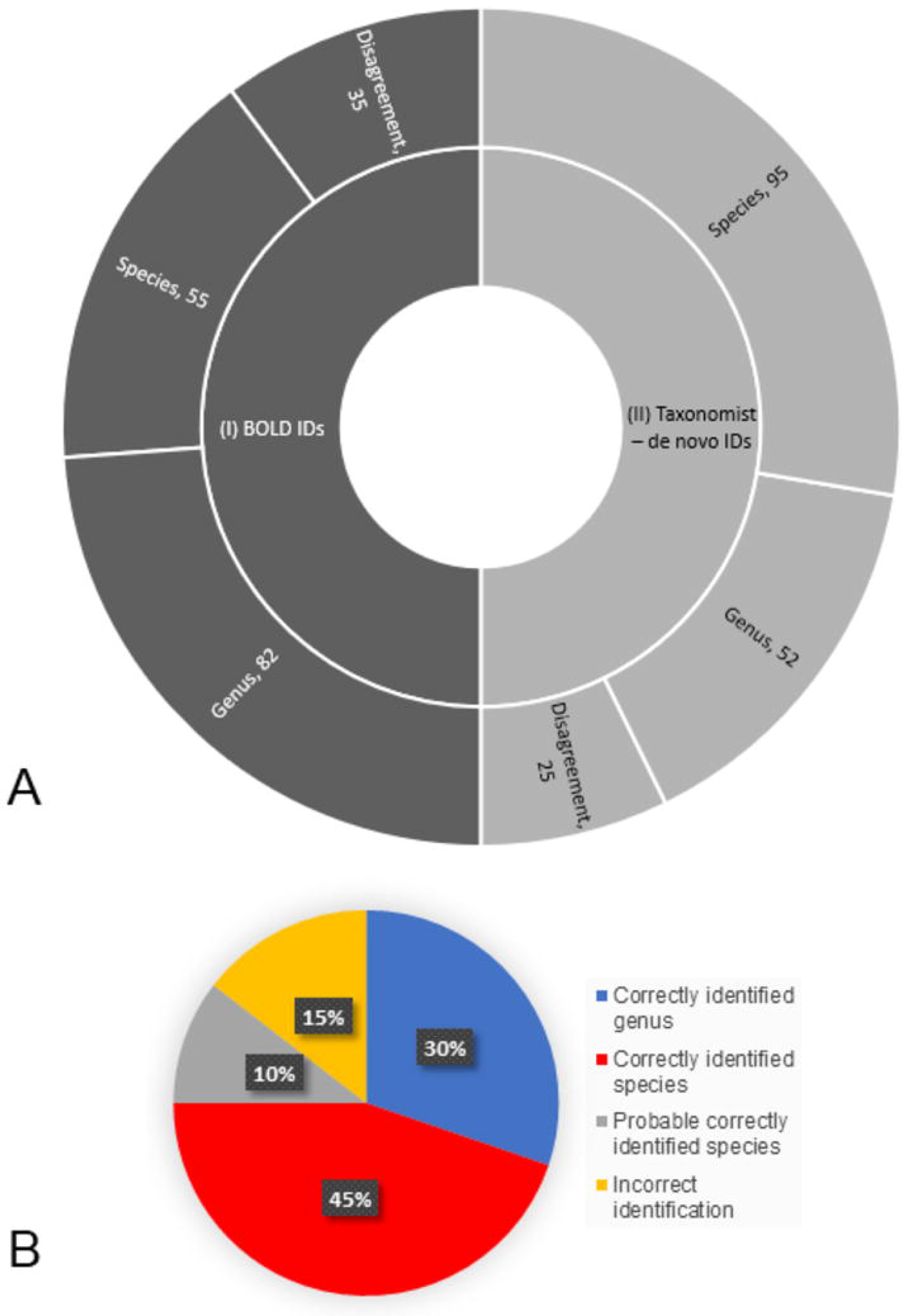
Results of the reanalysis performed by the taxonomist in the subfamily of dung beetles Scarabaeinae (Coleoptera: Scarabaeidae). A) Comparison between the identification (ID) matches before (I, BOLD IDs) and after (II, taxonomist – *de novo* IDs) reanalyzing the data. B) Percentage of the categories assigned by the taxonomist to the identifications suggested with BOLD’s search engine at 80%.

Then, we corroborated *de novo* the identification for these 172 records of dung beetle with a taxonomist (Fig 4B). The *de novo* specimens were assigned to the same 16 genera and 27 species (70 specimens), in addition to 31 morphospecies (genus and code “;H“, 97 specimens) (Table 4 and S4 Table). The new review resulted in 95 records (55%) correctly matched the taxonomist species identification or was deemed likely to be the species (Correctly identified species and Probable correctly identified species categories). At the genus level, 52 records (30%) correctly matched the taxonomist genus identification but either were not accurately matched with the species or the species identification was not provided (Correctly identified genus category). There were 25 records (15%) at the genus level which did not match the identification of the taxonomist and arguments against BOLD identification were provided by the taxonomist (Incorrect identification category) (S4 Table). Therefore, the records previously classified as 137 (80%) increased to 147 (85%) in agreement with the taxonomist and were reduced from 35 (20%) to 25 (15%) in disagreement based on the BOLD search engine at 80% (Table 3, Fig 4A).

**Table 4.** List of Scarabaeinae species identified by BOLD in agreement with the taxonomist’s *de novo* identification. Morphospecies with genus and code “H” are included [24]. Only categories for correct species and probably correct species are presented here.

| Process ID | BIN | Current ID | % Match | Overlap (bp) | Taxonomist's *de novo* ID | Match Species BOLD | Match BIN | Match notes |
| --- | --- | --- | --- | --- | --- | --- | --- | --- |
| CBIHM686-18 | BOLD:ACW4694 | *Ateuchus* | 93.4 | 568 | *Ateuchus aeneomicans* | *Ateuchus aeneomicans* | BOLD:AAW8107 | Correct species |
| CBIHI151-18 | BOLD:ADL3091 | *Canthon juvenicus* | 99.85 | 567 | *Canthon juvenicus* | *Canthon juvenicus* | BOLD:ADL3091 | Correct species |
| CBIHM706-18 | BOLD:ADL3091 | *Canthon juvenicus* | 99.85 | 567 | *Canthon juvenicus* | *Canthon juvenicus* | BOLD:ADL3091 | Correct species |
| CBIHM705-18 | BOLD:ADL3091 | *Canthon juvenicus* | 99.69 | 566 | *Canthon juvenicus* | *Canthon juvenicus* | BOLD:ADL3091 | Correct species |
| CBIHM633-18 |  | *Canthon* | 98.2 | 563 | *Sylvicanthon aequinoctialis* [30] | *Canthon aequinoctialis* | BOLD:ABA2649 | Correct species |
| CBIHM627-18 | BOLD:ABA2649 | *Canthon* | 98.01 | 561 | *Sylvicanthon aequinoctialis* | *Canthon aequinoctialis* | BOLD:ABA2649 | Correct species |
| CBIHM635-18 | BOLD:ABA2649 | *Canthon* | 97.55 | 570 | *Sylvicanthon aequinoctialis* | *Canthon aequinoctialis* | BOLD:ABA2649 | Correct species |
| CBIHM636-18 | BOLD:ABA2649 | *Canthon* | 97.55 | 570 | *Sylvicanthon aequinoctialis* | *Canthon aequinoctialis* | BOLD:ABA2649 | Correct species |
| CBIHM632-18 | BOLD:ABA2649 | *Canthon* | 97.25 | 570 | *Sylvicanthon aequinoctialis* | *Canthon aequinoctialis* | BOLD:ABA2649 | Correct species |
| CBIHM631-18 | BOLD:ABA2647 | *Canthon* | 100 | 570 | *Sylvicanthon aequinoctialis* | *Canthon aequinoctialis*ASolis1 | BOLD:ABA2647 | Correct species |
| CBIHM637-18 | BOLD:ABA2647 | *Canthon* | 100 | 570 | *Sylvicanthon aequinoctialis* | *Canthon aequinoctialis*ASolis1 | BOLD:ABA2647 | Correct species |
| CBIHM638-18 | BOLD:ABA2647 | *Canthon* | 100 | 570 | *Sylvicanthon aequinoctialis* | *Canthon aequinoctialis*ASolis1 | BOLD:ABA2647 | Correct species |
| CBIHM634-18 | BOLD:ABA2647 | *Canthon* | 99.85 | 570 | *Sylvicanthon aequinoctialis* | *Canthon aequinoctialis*ASolis1 | BOLD:ABA2647 | Correct species |
| CBIHM629-18 | BOLD:ABA2649 | *Canthon* | 98.17 | 570 | *Sylvicanthon aequinoctialis* | *Canthon aequinoctialis*ASolis3 | BOLD:ABA2649 | Correct species |
| CBIHM628-18 | BOLD:ABA2649 | *Canthon* | 98.08 | 570 | *Sylvicanthon aequinoctialis* | *Canthon aequinoctialis*ASolis3 | BOLD:ABA2649 | Correct species |
| CBIHM639-18 | BOLD:ABA2649 | *Canthon* | 97.86 | 570 | *Sylvicanthon aequinoctialis* | *Canthon aequinoctialis*ASolis3 | BOLD:ABA2649 | Correct species |
| CBIHM747-18 | BOLD:ACW3356 | *Canthon subhyalinus* | 100 | 570 | *Canthon subhyalinus* | *Canthon subhyalinus* | BOLD:ACW3356 | Correct species |
| CBIHM673-18 | BOLD:ACW3356 | *Canthon subhyalinus* | 100 | 570 | *Canthon subhyalinus* | *Canthon subhyalinus* | BOLD:ACW3356 | Correct species |
| CBIHM674-18 | BOLD:ACW3356 | *Canthon subhyalinus* | 100 | 570 | *Canthon subhyalinus* | *Canthon subhyalinus* | BOLD:ACW3356 | Correct species |
| CBIHM727-18 | BOLD:ACW3356 | *Canthon subhyalinus* | 100 | 570 | *Canthon subhyalinus* | *Canthon subhyalinus* | BOLD:ACW3356 | Correct species |
| CBIHM744-18 | BOLD:ACW3356 | *Canthon subhyalinus* | 100 | 567 | *Canthon subhyalinus* | *Canthon subhyalinus* | BOLD:ACW3356 | Correct species |
| CBIHM718-18 | BOLD:ACW3356 | *Canthon subhyalinus* | 99.24 | 570 | *Canthon subhyalinus* | *Canthon subhyalinus* | BOLD:ACW3356 | Correct species |
| CBIHM716-18 | BOLD:ACW3356 | *Canthon subhyalinus* | 98.93 | 570 | *Canthon subhyalinus* | *Canthon subhyalinus* | BOLD:ACW3356 | Correct species |
| CBIHM717-18 | BOLD:ACW3356 | *Canthon subhyalinus* | 98.93 | 570 | *Canthon subhyalinus* | *Canthon subhyalinus* | BOLD:ACW3356 | Correct species |
| CBIHM741-18 | BOLD:ACW3356 | *Canthon* | 100 | 570 | *Canthon subhyalinus* | *Canthon subhyalinus* | BOLD:ACW3356 | Correct species |
| CBIHI156-18 | BOLD:ADO2303 | *Canthon triangularis* | 100 | 570 | *Canthon triangularis* | *Canthon triangularis* | BOLD:ADO2303 | Correct species |
| CBIHI157-18 | BOLD:ADO2303 | *Canthon triangularis* | 100 | 570 | *Canthon triangularis* | *Canthon triangularis* | BOLD:ADO2303 | Correct species |
| CBIHM659-18 | BOLD:ADL4806 | *Coprophanaeus corythus* | 97.24 | 540 | *Coprophanaeus corythus* | *Coprophanaeus corythus* | BOLD:ACW3300 | Correct species |
| CBIHM687-18 | BOLD:ADL4607 | *Coprophanaeus gamezi* | 94.65 | 570 | *Coprophanaeus gamezi* | *Coprophanaeus gamezi* | BOLD:ACW3623 | Correct species |
| CBIHI019-17 | BOLD:ADJ7028 | *Deltochilum mexicanum* | 95.26 | 570 | *Deltochilum mexicanum* | *Deltochilum mexicanum* | BOLD:AAP9113 | Correct species |
| CBIHI159-18 | BOLD:ACW3574 | *Dichotomius* | 95.61 | 570 | *Dichotomius agenor* | *Dichotomius agenor* | BOLD:ABA2814 | Correct species |
| CBIHM648-18 | BOLD:ACW3574 | *Dichotomius* | 95.61 | 570 | *Dichotomius agenor* | *Dichotomius agenor* | BOLD:ABA2814 | Correct species |
| CBIHM667-18 | BOLD:ACW3574 | *Dichotomius* | 95.36 | 570 | *Dichotomius agenor* | *Dichotomius agenor* | BOLD:ABA2814 | Correct species |
| CBIHM666-18 | BOLD:ACW3574 | *Dichotomius* | 95.28 | 570 | *Dichotomius agenor* | *Dichotomius agenor* | BOLD:ABA2814 | Correct species |
| CBIHM646-18 | BOLD:ACW3574 | *Dichotomius* | 95.23 | 570 | *Dichotomius agenor* | *Dichotomius agenor* | BOLD:ABA2814 | Correct species |
| CBIHM647-18 | BOLD:ACW3574 | *Dichotomius* | 95.21 | 567 | *Dichotomius agenor* | *Dichotomius agenor* | BOLD:ABA2814 | Correct species |
| CBIHM645-18 | BOLD:ACW3574 | *Dichotomius* | 95.12 | 570 | *Dichotomius agenor* | *Dichotomius agenor* | BOLD:ABA2814 | Correct species |
| CBIHM665-18 | BOLD:ACW3574 | *Dichotomius* | 95.12 | 570 | *Dichotomius agenor* | *Dichotomius agenor* | BOLD:ABA2814 | Correct species |
| CBIHI144-18 | BOLD:AAC9021 | *Digitonthophagus gazella* | 100 | 570 | *Digitonthophagus gazella* | *Digitonthophagus gazella* | BOLD:AAC9021 | Correct species |
| CBIHM640-18 | BOLD:ADL2649 | *Eurysternus caribaeus* | 96.48 | 570 | *Eurysternus caribaeus* | *Eurysternus caribaeus* | BOLD:ABA7207 | Correct species |
| CBIHI145-18 | BOLD:ACW4232 | *Eurysternus caribaeus* | 100 | 570 | *Eurysternus caribaeus* | *Eurysternus caribaeus* | BOLD:ACW4232 | Correct species |
| CBIHI146-18 | BOLD:ACW4232 | *Eurysternus caribaeus* | 100 | 570 | *Eurysternus caribaeus* | *Eurysternus caribaeus* | BOLD:ACW4232 | Correct species |
| CBIHI147-18 | BOLD:ACW4232 | *Eurysternus caribaeus* | 100 | 570 | *Eurysternus caribaeus* | *Eurysternus caribaeus* | BOLD:ACW4232 | Correct species |
| CBIHI023-17 | BOLD:ADJ7677 | *Eurysternus foedus* | 95.57 | 570 | *Eurysternus foedus* | *Eurysternus foedus* | BOLD:ABU6799 | Correct species |
| CBIHI030-17 | BOLD:AAX0271 | *Onthophagus acuminatus* | 100 | 570 | *Onthophagus acuminatus* | *Onthophagus acuminatus*AS1 | BOLD:AAX0271 | Correct species |
| CBIHM653-18 | BOLD:ADL2546 | *Eurysternus mexicanus* | 100 | 570 | *Eurysternus mexicanus* | *Eurysternus mexicanus* | BOLD:ADL2546 | Correct species |
| CBIHM654-18 | BOLD:ADL2546 | *Eurysternus mexicanus* | 100 | 570 | *Eurysternus mexicanus* | *Eurysternus mexicanus* | BOLD:ADL2546 | Correct species |
| CBIHM655-18 | BOLD:ADL2546 | *Eurysternus mexicanus* | 100 | 570 | *Eurysternus mexicanus* | *Eurysternus mexicanus* | BOLD:ADL2546 | Correct species |
| CBIHM656-18 | BOLD:ADL2546 | *Eurysternus mexicanus* | 100 | 570 | *Eurysternus mexicanus* | *Eurysternus mexicanus* | BOLD:ADL2546 | Correct species |
| CBIHM660-18 | BOLD:ADL2546 | *Eurysternus mexicanus* | 100 | 570 | *Eurysternus mexicanus* | *Eurysternus mexicanus* | BOLD:ADL2546 | Correct species |
| CBIHM661-18 | BOLD:ADL2546 | *Eurysternus mexicanus* | 100 | 570 | *Eurysternus mexicanus* | *Eurysternus mexicanus* | BOLD:ADL2546 | Correct species |
| CBIHM662-18 | BOLD:ADL2546 | *Eurysternus mexicanus* | 100 | 570 | *Eurysternus mexicanus* | *Eurysternus mexicanus* | BOLD:ADL2546 | Correct species |
| CBIHM657-18 | BOLD:ADL2546 | *Eurysternus mexicanus* | 99.83 | 570 | *Eurysternus mexicanus* | *Eurysternus mexicanus* | BOLD:ADL2546 | Correct species |
| CBIHM698-18 | BOLD:ADL4924 | *Malagoniella astyanax* | 94.34 | 570 | *Malagoniella astyanax* | *Malagoniella astyanax* | BOLD:ABA3615 | Correct species |
| CBIHM701-18 | BOLD:ACW3925 | *Ontherus appendiculatus* | 99 | 558 | *Ontherus appendiculatus* | *Ontherus appendiculatus* | BOLD:ACW3925 | Correct species |
| CBIHM703-18 | BOLD:ACW3925 | *Ontherus appendiculatus* | 99 | 570 | *Ontherus appendiculatus* | *Ontherus appendiculatus* | BOLD:ACW3925 | Correct species |
| CBIHM699-18 | BOLD:ACW3925 | *Ontherus appendiculatus* | 98.95 | 570 | *Ontherus appendiculatus* | *Ontherus appendiculatus* | BOLD:ACW3925 | Correct species |
| CBIHM702-18 | BOLD:ACW3925 | *Ontherus appendiculatus* | 98.5 | 558 | *Ontherus appendiculatus* | *Ontherus appendiculatus* | BOLD:ACW3925 | Correct species |
| CBIHI160-18 |  | *Onthophagus curvicornis* | 99.82 | 552 | *Onthophagus curvicornis* | *Onthophagus curvicornis* | BOLD:ADM9496 | Correct species |
| CBIHI142-18 | BOLD:ADM9497 | *Onthophagus curvicornis* | 98.17 | 570 | *Onthophagus curvicornis* | *Onthophagus curvicornis* | BOLD:ADM9497 | Correct species |
| CBIHI161-18 | BOLD:ADM9497 | *Onthophagus curvicornis* | 98.17 | 570 | *Onthophagus curvicornis* | *Onthophagus curvicornis* | BOLD:ADM9497 | Correct species |
| CBIHI162-18 | BOLD:ADM9496 | *Onthophagus curvicornis* | 99.82 | 567 | *Onthophagus curvicornis* | *Onthophagus curvicornis* |  | Correct species |
| CBIHM719-18 | BOLD:ADL5334 | *Onthophagus lebasii* | 99.84 | 569 | *Onthophagus lebasii* | *Onthophagus lebasii* | BOLD:ADL5334 | Correct species |
| CBIHM735-18 | BOLD:ADL5334 | *Onthophagus lebasii* | 99.84 | 567 | *Onthophagus lebasii* | *Onthophagus lebasii* | BOLD:ADL5334 | Correct species |
| CBIHM726-18 | BOLD:ADL5334 | *Onthophagus lebasii* | 99.35 | 567 | *Onthophagus lebasii* | *Onthophagus lebasii* | BOLD:ADL5334 | Correct species |
| CBIHM721-18 | BOLD:ADL5334 | *Onthophagus lebasii* | 98.61 | 566 | *Onthophagus lebasii* | *Onthophagus lebasii* | BOLD:ADL5334 | Correct species |
| CBIHI152-18 | BOLD:ADM9873 | *Phanaeus bispinus* | 96.94 | 570 | *Phanaeus bispinus* | *Phanaeus bispinus* | BOLD:ACW4591 | Correct species |
| CBIHI153-18 | BOLD:ADN3451 | *Phanaeus haroldi* | 98.62 | 570 | *Phanaeus haroldi* | *Phanaeus haroldi* | BOLD:ADN3451 | Correct species |
| CBIHI154-18 | BOLD:ADN3451 | *Phanaeus haroldi* | 98.62 | 570 | *Phanaeus haroldi* | *Phanaeus haroldi* | BOLD:ADN3451 | Correct species |
| CBIHM712-18 | BOLD:ACW3786 | *Sulcophanaeus leander* | 99.83 | 570 | *Sulcophanaeus leander* | *Sulcophanaeus leander* | BOLD:ACW3786 | Correct species |
| CBIHM713-18 | BOLD:ACW3786 | *Sulcophanaeus leander* | 99.68 | 570 | *Sulcophanaeus leander* | *Sulcophanaeus leander* | BOLD:ACW3786 | Correct species |
| CBIHM711-18 | BOLD:ACW3786 | *Sulcophanaeus leander* | 99.66 | 570 | *Sulcophanaeus leander* | *Sulcophanaeus leander* | BOLD:ACW3786 | Correct species |
| CBIHM729-18 | BOLD:ADL4592 | *Uroxys micros* | 100 | 504 | *Uroxys micros* | *Uroxys micros* | BOLD:ADL4592 | Correct species |
| CBIHM731-18 | BOLD:ADL4592 | *Uroxys micros* | 100 | 501 | *Uroxys micros* | *Uroxys micros* | BOLD:ADL4592 | Correct species |
| CBIHM740-18 | BOLD:ADL4592 | *Uroxys micros* | 100 | 504 | *Uroxys micros* | *Uroxys micros* | BOLD:ADL4592 | Correct species |
| CBIHM746-18 | BOLD:ADL4592 | *Uroxys micros* | 99.83 | 513 | *Uroxys micros* | *Uroxys micros* | BOLD:ADL4592 | Correct species |
| CBIHM739-18 | BOLD:ADL4592 | *Uroxys micros* | 99.16 | 494 | *Uroxys micros* | *Uroxys micros* | BOLD:ADL4592 | Correct species |
| CBIHI025-17 | BOLD:ADK7115 | *Canthidium* | 95.39 | 567 | *Canthidium* sp. 21H | *Canthidium centrale* | BOLD:ABU9359 | Probably correct species |
| CBIHM650-18 | BOLD:ACW2842 | *Canthon* | 94.97 | 567 | *Canthon* sp. 06H | *Canthon cyanellus* | BOLD:ADN2169 | Probably correct species |
| CBIHM651-18 | BOLD:ACW2842 | *Canthon* | 94.97 | 567 | *Canthon* sp. 06H | *Canthon cyanellus* | BOLD:ADN2169 | Probably correct species |
| CBIHM663-18 | BOLD:ACW2842 | *Canthon* | 94.9 | 570 | *Canthon* sp. 06H | *Canthon cyanellus* | BOLD:ADN2169 | Probably correct species |
| CBIHM664-18 | BOLD:ACW2842 | *Canthon* | 94.9 | 570 | *Canthon* sp. 06H | *Canthon cyanellus* | BOLD:ADN2169 | Probably correct species |
| CBIHM649-18 | BOLD:ACW2842 | *Canthon* | 94.85 | 567 | *Canthon* sp. 06H | *Canthon cyanellus* | BOLD:ADN2169 | Probably correct species |
| CBIHM742-18 | BOLD:ACW4541 | *Canthon* | 96.83 | 570 | *Canthon* sp. 09H | *Canthon caelius* | BOLD:ABU7368 | Probably correct species |
| CBIHM668-18 | BOLD:ACW4541 | *Canthon* | 96.62 | 567 | *Canthon* sp. 09H | *Canthon caelius* | BOLD:ABU7368 | Probably correct species |
| CBIHM669-18 | BOLD:ACW4541 | *Canthon* | 96.62 | 567 | *Canthon* sp. 09H | *Canthon caelius* | BOLD:ABU7368 | Probably correct species |
| CBIHM670-18 | BOLD:ACW4541 | *Canthon* | 96.62 | 567 | *Canthon* sp. 09H | *Canthon caelius* | BOLD:ABU7368 | Probably correct species |
| CBIHM730-18 | BOLD:ACW4541 | *Canthon* | 96.62 | 567 | *Canthon* sp. 09H | *Canthon caelius* | BOLD:ABU7368 | Probably correct species |
| CBIHM732-18 | BOLD:ACW4541 | *Canthon* | 96.6 | 570 | *Canthon* sp. 09H | *Canthon caelius* | BOLD:ABU7368 | Probably correct species |
| CBIHM820-20 |  | *Canthon* | 95.11 | 408 | *Canthon* sp. 09H | *Canthon caelius* | BOLD:ABU7368 | Probably correct species |
| CBIHM683-18 | BOLD:ADL3950 | *Canthon* | 95.01 | 570 | *Canthon* sp. 23H | *Canthon bispinus* | BOLD:ADR8659 | Probably correct species |
| CBIHM684-18 | BOLD:ADL3950 | *Canthon* | 95.01 | 570 | *Canthon* sp. 23H | *Canthon bispinus* | BOLD:ADR8659 | Probably correct species |
| CBIHI148-18 | BOLD:AAX0252 | *Onthophagus* | 98.04 | 570 | *Onthophagus* sp. 07H | *Onthophagus haematopus* | BOLD:AAX0252 | Probably correct species |
| CBIHI150-18 | BOLD:AAX0252 | *Onthophagus* | 98.04 | 570 | *Onthophagus* sp. 07H | *Onthophagus haematopus* | BOLD:AAX0252 | Probably correct species |
| CBIHI149-18 | BOLD:AAX0252 | *Onthophagus* | 97.88 | 570 | *Onthophagus* sp. 07H | *Onthophagus haematopus* | BOLD:AAX0252 | Probably correct species |

Correctly identified species had a percentage match range in BOLD from 93.4 to 100 (Fig 5). Probable correctly identified species ranged from 94.85 to 98.04. Correctly identified genus range was between 87.7 and 100%. Incorrect identification percentages range from 83.59 to 99.83. The categories were significantly different (Kruskal-Wallis H = 73.4, P < 0.001). The correctly identified species (P < 0.001), probable correctly identified species (P < 0.05) and correctly identified genus (P < 0.05) categories were different from the incorrect identifications.

**Fig 5.**
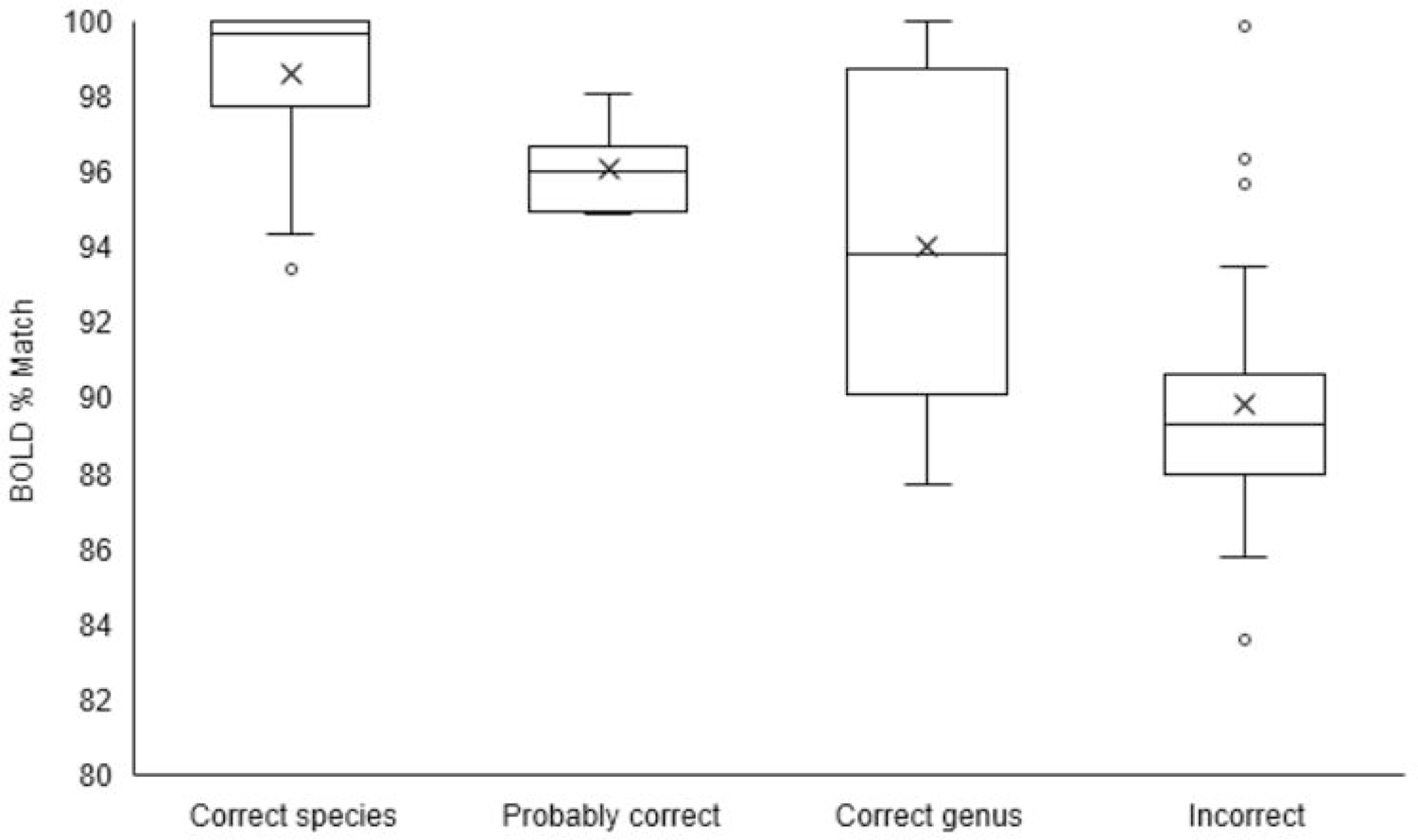
Boxplot of Scarabaeinae case of study showing the range of BOLD percentage matches for the categories: Correctly Identified Species, Probable Correctly Identified Species, Correctly Identified Genus, and Incorrect Identification. The median is represented by a line, the mean is represented by an X markers and outliers are the white dots.

## Discussion

BOLD systems outperformed BLAST providing more accurate insect identifications. This overall pattern was found for all taxonomic levels evaluated (species, genus and family). Although differences were found between taxa, BOLD consistently provided higher identifications for the order Coleoptera at all taxonomic levels and for Lepidoptera at genus and species level. For instance, Hymenoptera and Odonata performed similarly for both repositories at all taxonomic levels, alike most of the other orders at the family level when analyzed separately. These results are partially in disagreement with Meiklejohn et al [21] which reported no significant differences between BOLD and GenBank for insect samples identification at genus and species level. A further review of their data showed that correct identifications from both platforms were closer than estimated previously [22]. Although we report different results in the performance of BOLD compared to GenBank in this study, a key factor to consider is the magnitude of insect data involved. Meiklejohn et al [21], who studied 17 specimens in 12 orders at genus and species level using COI whereas we included 1160 specimens in eight orders at family, genus and species level. BOLD Systems is a platform developed with the intention of providing species identification and in that sense does not perform equally to GenBank, but both are the main public repositories often used to provide identifications. Often BOLD is considered to contain data more curated, but GenBank data quality is good overall [22,31]. Differences in identifications of taxa can be considered due to differential taxonomic curation of the databases and the lack of barcodes of reference from Colombia. However, we could not identify a clear pattern with the lack of references. The most represented groups were Coleoptera and Lepidoptera. The group that performed the best with identifications was Coleoptera and it was also the order that provided the most unique BINS, in contrast with Lepidoptera that provided the least unique BINS also outperformed GenBank at the species and genus level, but no differences were found at family.

A great potential of Colombian insect biodiversity was found from BINs counting with 70% of the sequences assigned to a unique BIN. Each BIN represents a cluster of species or an operational taxonomic unit (OTU) that is compared against the database assigning the barcodes to known clusters (BIN) or creating a new cluster (unique BIN) [18]. Unique BINs were found across all the orders of insects and Coleoptera provided the highest count of new BINs followed by Diptera and Hymenoptera, some of the hyperdiverse orders of insects. However, Lepidoptera was the order with highest non-unique BINs. This finding is likely attributable to a great effort of DNA sampling directed to Lepidoptera that displays 4848 barcodes from the country with 1474 BINs [32]; in comparison to Coleoptera with 318 sequences in 82 BINs, Diptera 2646 with 339 BINs and Hymenoptera 2179 with 240 BINs [32].

Furthermore, our discovery of 500 unique BINs from our dataset of 1088 sequences showcases the need for further sampling and identification in order to add more species into these databases and increase the accuracy of matches. However, this would require the implementation of a systematic method of collecting samples across Colombia, a process that could take a considerable length of time and amount of labor. Of our seven collecting methods used in this study, the malaise traps accounted for 62.6%, and we would suggest a targeted approach by placing these traps in strategic areas across Colombia. We expect that a larger sample will continue providing a higher number of unique BINs, but also, we see a necessary taxonomic effort to match all of this genetic diversity with the insect species already described from Colombia. Nevertheless, the implementation of DNA barcodes in the country will bring an opportunity to recognize any hidden diversity. As it has been suggested that BINs underestimate by a 10% the species due to lack of power within close species [11,33,34]. Although Meier et al [35] suggested awareness on the use of BINS as identification tool specially when use for species descriptions. Therefore, an integrative approach with morphology and genetic diversity can boost the recognition of species.

A closer look to the subfamily Scarabaeinae displayed a great performance of identification at different levels. The percentage of similarity for a match (BOLD match percentage) to identify species correctly in Scarabaeinae was in a wide range between 93 to a 100%. Although correct matches were statistically significant the range was not perfect, as shown at least three specimens were incorrectly identified with values higher than 94%. These outliers do not reduce the potential of genetic identification tools. The high percentages could be due to specimen mix-up. Overall, the identification success rate of Scarabaeinae was 85% (30% towards genera and 55% towards species) when combining the barcode results with a taxonomist effort (Table 3, Fig 4A). The species identification improved from 78% to 96% (or in the overall from 30% to 55%) with this taxonomist *de novo* identifications (Table 3, Fig 4A). Other studies of identification of beetles using barcodes found species matches as high as 92.1% of their samples when considered traditionally identified species and BINS and as high as 98.3% of the samples when haplotypes were considered [36]. In comparison, our identification percentage was lower with previous studies. These can be due to the lack of sequence records of the species in the database. Pentisaari et al [36] suggested identification failures due to difficulty with morphological characters and controversial taxonomic status of species. Taxonomic input is crucial for the function of this tool, also if the species is not represented in the database there is no possible match to occur. In fact, a few suggestions provided by BOLD for incorrect species identifications were for close related species or species within their group of species. For example, *Dichotomius andresi* and *Dichotomius protectus* were matched with *Dichotomius satanas*, a morphologically related species in the “*Dichotomius satanas* species group”. Similarly, *Eurysternus hypocrita* was identified as *Eurysternus olivaceus*, a species in the “*Eurysternus velutinus* species group". The specimen identified as *Phanaeus pyrois* was matched as *Phanaeus malyi*, a species that was long considered synonymous with *P. pyrois*. Currently, *P. malyi* has been revalidated and registered in Colombia without a specific location [37].

The distribution range of species could be an additional factor as observed with the dung beetles data where BOLD identify all species correctly in six genera (*Coprophanaeus, Digitonthophagus, Malagoniella, Onthophagus, Phanaeus* and *Sulcophanaeus*), eight genera partially correct with some of the species identified correctly but not all (*Ateuchus, Canthidium, Canthon, Deltochilum, Dichotomius, Eurysternus, Ontherus* and *Uroxys*) and failed completely for the genera *Pseudocanthon* (1 specimen) and *Scybalocanthon* (6 specimens). The correctly identified species were obtained for species with a wide distribution and shared between South America and Central America, such as *Canthon juvencus, Canthon subhyalinus, Eurysternus caribaeus, Eurysternus foedus, Eurysternus mexicanus Onthophagus acuminatus, Onthophagus lebasi*, and *Uroxys micros*, most likely due to representativeness of COI sequences of these species from Costa Rica [38]. These species are found across countries or ecosystems (e. g. dry and humid forests, natural and disturbed ecosystems). In contrast, the identification was incorrect for some species with restricted distributions in Colombia such as *Canthon arcabuquensis, Dichotomius andresi, Uroxys cuprescens*, and *Deltochilum susanae*, the latest was first identified as the morphospecies *Deltochilum* sp. 12H [39] and later described as a new species [40] or species with Andean distributions as in *Dichotomius protectus, Eurysternus marmoreus* and *Ontherus brevicollis* [41–46]. In the case of Scarabaeinae dung beetles, the barcode results helped to confirm at least five species that were assigned as morphospecies in the dung beetles of Colombia “Reference Collection” hosted at of IAVH Entomological Collection. The diversity of dung beetle’s subfamily Scarabaeinae in Colombia exceeds by a large number of species that of neighboring countries such as Panama, Ecuador or Brazil. In general, a defined and well-identified species in Colombia has other species, morphologically similar and taxonomically closer, that have been assigned as morphospecies. This is why the Reference Collection was created, which so far houses about 85% percent of species of Colombia as morphospecies. We see the barcode as a fundamental integrative taxonomy tool, which will help to outline these morphospecies and define which of those are new species for science, and to improve the final taxonomic list of this important group of beetles. Overall, there are still challenges to overcome and more research is recommended in this topic. The reference databases keep improving and updating their resources and tools for users [20,31]. BOLD provided reliable identifications for the tested groups of insects overcoming challenges such as a small genetic reference of insects for a highly diverse country.

## Acknowledgements

We are grateful to our funding sources: A Minciencias postdoctoral fellowship Fondo Nacional de Financiamiento para la Ciencia, la Tecnología y la Innovación “Francisco José de Caldas” call No. 848-2019 and Humboldt Institute (awarded to NBB), Colombia BIO Agreement # FP 44842-109-2016 (IAvH No. 16-062) between Colciencias (now Minciencias) and Humboldt Institute, and Santander BIO inter-agency Agreement #2243 (IAvH No. 17-199) with the General System of Royalties of the Santander government, Colombia’s National Planning Department (BPIN 2017000100046), Humboldt Institute and Industrial University of Santander (BPIN 2017000100046). We would like to thank the following researchers for their fieldwork efforts collecting the samples and/or for identifications of samples: H Arenas, C Beltran, C Bota, J Cardenas, C Castro, A Clavijo, Y Correa, A Diaz, D Espitia, G Fagua, A González, D Hincapie, A Lopera, J Mendivil, R Morales, EE Palacio, A Parrales, P Pulido, JD Suaza, EA Tenorio, E Torres, X. Villalba. We want to thank to the valuable Entomological collections where these samples were deposited: the Entomological and Tissue collection of the Institute Alexander von Humboldt (IAvH-E and IAvH-CT), Museo Francisco Luis Gallego (MEFLG) at Universidad Nacional de Colombia, sede Medellín, Museo Javeriano de Historia Natural (MPUJ) at Pontificia Universidad Javeriana and Colección Entomológica Universidad de Antioquia (CEUA). Thank you to the IAvH Conservation Genetic lab members: P Montoya, P Pulido, N Franco, V Sandoval, C Quiroga, and Y Castañeda for their valuable comments and improvements to the manuscript.

## Author Contributions

**Conceptualization:** Nathalie Baena-Bejarano, Mailyn Gonzalez.

**Data curation:** Nathalie Baena-Bejarano, Catalina Reina, Diego E. Martínez-Revelo.

**Formal analysis:** Nathalie Baena-Bejarano.

**Investigation:** Nathalie Baena-Bejarano, Diego E. Martínez-Revelo, Claudia A. Medina, Jhon Cesar Neita, Catalina Reina, Eduardo Tovar, Sandra Uribe, Mailyn Gonzalez.

**Methodology:** Nathalie Baena-Bejarano, Catalina Reina, Diego E. Martínez-Revelo, Claudia A. Medina, Eduardo Tovar, Mailyn Gonzalez.

**Project administration:** Nathalie Baena-Bejarano, Mailyn Gonzalez, Jhon Cesar Neita.

**Supervision:** Mailyn Gonzalez, Jhon Cesar Neita, Nathalie Baena-Bejarano.

**Resources:** Mailyn Gonzalez, Claudia A. Medina, Jhon Cesar Neita, Eduardo Tovar, Diego E. Martínez-Revelo.

**Writing – original draft Preparation:** Nathalie Baena-Bejarano.

**Writing – review & editing:** Nathalie Baena-Bejarano, Diego E. Martínez-Revelo, Claudia A. Medina, Catalina Reina, Eduardo Tovar, Jhon Cesar Neita, Mailyn Gonzalez, Sandra Uribe.

## Supporting information captions

**S1 Table. Project dataset with the insect records**.

**S2 Table. Records of insects by orders with matches from BOLD Batch ID engine and BLAST (GenBank)**.

**S3 Table. Chi-squared tests**. A) Dataset analyses including BOLD’s first suggestion with the highest percentage match and base pair overlap. This dataset includes matches to sequences on our project. B) Alternative dataset analyses including next higher percentage match and base pair overlap. This excludes matches to sequences from our project. *Is next to orders where Chi-square was significant. **Is next to orders where Chi-square analyzed with Bonferroni correction was significant.

**S4 Table. List of Scarabaeinae in other categories as determined by the taxonomist**.

